# Cytosolic and mitochondrial translation elongation are coordinated through the molecular chaperone TRAP1 for the synthesis and import of mitochondrial proteins

**DOI:** 10.1101/2023.01.19.524708

**Authors:** Rosario Avolio, Ilenia Agliarulo, Daniela Criscuolo, Daniela Sarnataro, Margherita Auriemma, Sara Pennacchio, Giovanni Calice, Martin Y. Ng, Carlotta Giorgi, Paolo Pinton, Barry Cooperman, Matteo Landriscina, Franca Esposito, Danilo Swann Matassa

**Author notes:** The authors wish it to be known that, in their opinion, the first 2 authors should be regarded as joint First Authors. Correspondence may also be addressed to: Franca Esposito.

## Abstract

A complex interplay between mRNA translation and cellular respiration has been recently unveiled, but its regulation in humans is poorly characterized in either health or disease. Cancer cells radically reshape both biosynthetic and bioenergetic pathways to sustain their aberrant growth rates. In this regard, we have shown that the molecular chaperone TRAP1 not only regulates the activity of respiratory complexes, behaving alternatively as an oncogene or a tumor suppressor, but also plays a concomitant moonlighting function in mRNA translation regulation. Herein we identify the molecular mechanisms involved, demonstrating that TRAP1: i) binds both mitochondrial and cytosolic ribosomes as well as translation elongation factors, ii) slows down translation elongation rate, and iii) favors localized translation in the proximity of mitochondria. We also provide evidence that TRAP1 is coexpressed in human tissues with the mitochondrial translational machinery, which is responsible for the synthesis of respiratory complex proteins. Altogether, our results show an unprecedented level of complexity in the regulation of cancer cell metabolism, strongly suggesting the existence of a tight feedback loop between protein synthesis and energy metabolism, based on the demonstration that a single molecular chaperone plays a role in both mitochondrial and cytosolic translation, as well as in mitochondrial respiration.

## INTRODUCTION

The flow of genetic information from DNA to protein entails multiple highly regulated steps from mRNA transcription, processing, and export, to translation into proteins with subsequent folding, post-translational modification, and, eventually, degradation. Moreover, proteins imported or integrated into organelles are targeted to their destination on the basis of signals in the peptide sequence or even upstream through their mRNA localization (1). In the case of mitochondria, the coupling between protein import machinery, assembly and activity of respiratory-chain complexes and regulation of mitochondrial protein synthesis adds a further level of complexity. In yeast, recent reports have shown that mitochondrial and cytosolic translation are rapidly, dynamically and synchronously regulated. Thus, the nuclear genome coordinates mitochondrial and cytosolic translation to orchestrate the timely synthesis of oxidative phosphorylation complexes, representing a previously underestimated regulatory layer shaping the mitochondrial proteome (2). Accordingly, the mitochondrial protein import machinery has multiple connections to the respiratory chain (3) and, in turn, the mitochondrial import receptor TOM20 mediates localization to the mitochondria outer membrane of mRNAs encoding mitochondrial proteins in a translation-dependent manner (4). At present, such a level of complexity is poorly explored in higher organisms, either in health or disease. Contrary to conventional wisdom, nowadays outdated, functional mitochondria are essential for cancer cells (5). Although mutations in mitochondrial genes are common in cancer cells, they do not cause the inactivation of mitochondrial energy metabolism, but rather alter the mitochondrial bioenergetic and biosynthetic status (6). Then communication with the nucleus through a mitochondrial ‘retrograde signaling’ leads to modulation of signal transduction pathways, transcriptional circuits and chromatin structure to meet the mitochondrial and nuclear requirements of the cancer cell. Among the numerous players involved in this process, the molecular chaperone TRAP1 has emerged in the last decade as a critical regulator of metabolic remodeling in cancer cells (7). TRAP1 was initially described as a chaperone for the retinoblastoma protein during mitosis and after heat shock (8), as a TNF-Receptor associated protein (9), and as a factor stabilizing CypD, which prevents permeability transition pore opening and thus apoptosis (10). However, in the last few years TRAP1 has emerged as a modulator of mitochondrial respiration, both through direct binding to respiratory complex II (11,12) and III (13), and indirect modulation of complex IV activity (14). The regulation of cancer cell metabolism by TRAP1 appears to have contextual effects on cancer onset and progression, thus favoring the oncogenic phenotype in glycolytic tumors (15), while being negatively selected in tumors mostly relying on oxidative metabolism (16). Conversely, TRAP1 also participates in biosynthetic pathways, through the coupled regulation of protein synthesis and degradation (17) and the remodeling of cholesterol homeostasis in ovarian cancer cells (18). Indeed, TRAP1 is also partly localized on the endoplasmic reticulum membrane (19), facing the cytosol, where it binds translation factors (17) and ribosomes (20).

Herein, we further characterize molecular mechanisms involved in TRAP1 translational regulation, showing for the first time that it participates in protein synthesis control both inside and outside mitochondria, by binding ribosomes and translation elongation factors, and contributing to co-translational protein localization to mitochondria. Herein the first demonstration is provided that human mitochondrial and cytosolic translation share a common regulatory molecular chaperone, thus suggesting possible mechanisms of coordinated regulations and feedback loops between protein synthesis and energy metabolism.

## MATERIAL AND METHODS

### Cell cultures

Human HCT116 colon carcinoma cells and human cervical carcinoma HeLa cells were purchased from American Type Culture Collection (ATCC) and cultured in McCoy’s 5A medium (HCT116) and DMEM (HeLa). Both culturing media contain 10% fetal bovine serum, 1.5 mmol/L glutamine. The authenticity of the cell lines was verified by STR profiling, in accordance with ATCC product description. HeLa Flp In Trex (FITR) cell line were kindly provided by Dr. Matthias Gromeier (Duke University Medical Center, Durham, USA). Generation of the HeLa Flp In Trex stable cell lines expressing the eGFP- or Flag-fusion proteins or the short hairpin RNA, was performed as described in the manufacturer’s protocol (Flp In Trex, Invitrogen). HeLa Flp In Trex cells were cultured in DMEM supplemented with 10% fetal bovine serum, 1.5 mmol/L glutamine, and appropriate selective antibiotics. Addition of tetracycline induces proteins as described previously (21).

### Plasmid generation and transfection procedures

For TRAP1-eGFP and TRAP1-Flag plasmids generation, HeLa cDNA library, eGFP and Flag plasmid were used as templates for fusion PCR. The resulting chimeric cDNAs were cloned into pCDNA5/FRT/TO. pFRT-U6tetO is a kind gift from prof. John J Rossi. Inducible short hairpin (sh) RNAs were generated as described in (22) (using BglII/KpnI as restriction sites). Short hairpin sequences used were: GFP=agatctGCACAAGCTGGAGTACAACTACCTGACCCATAGTTGTACTCCAGCTTGTGCTTTTTgg tacc; TRAP1=agatctGCCCGGTCCCTGTACTCAGAAACCTGACCCATTTCTGAGTACAGGGACCGGGCTTTTTggtacc.

### Polysome profiling

3×10 cm plates of cells were incubated 15 min at 37°C with fresh medium supplemented with 100 μg/mL of cycloheximide (Sigma). Where indicated cells were treated with 2 μg/mL of harringtonine or 100 μg/mL of puromycin. Cells were then washed with ice-cold PBS supplemented with 100 μg/mL cycloheximide and resuspended in 1 mL lysis buffer (10 mM Tris-HCl pH7.4, 100 mM KCl, 10 mM MgCl2, 1% Triton-X100, 1 mM DTT, 10 U/mL RNAseOUT (Invitrogen), 100 μg/mL of cycloheximide). After 5 min of incubation on ice, cell lysate was centrifuged for 10 min at 14000 rpm at 4°C. Where indicated cell extract was treated with 30 mM EDTA to disassemble polysomes, as a negative control. The supernatant was collected, and the absorbance was measured at 260 nm with the NanoDrop. Eight A260 units were loaded onto a 10-60% sucrose gradient obtained by adding 6 mL of 10% sucrose over a layer of 6 mL of 60% sucrose prepared in lysis buffer without Triton and containing 0.5 mM DTT, in a 12-mL tube (Polyallomer; Beckman Coulter). Gradients were prepared using a gradient maker (Gradient Master, Biocomp). Polysomes were separated by centrifugation at 35000 rpm for 2 hrs using a Beckmann SW41 rotor. Twelve fractions of 920 μL were collected while polysomes were monitored by following the absorbance at 254 nm. Total protein was retrieved by 20% trichloroacetic acid (TCA) precipitation performed overnight, washed with Acetone and analyzed by SDS-PAGE followed by Western blot. RNA extraction was performed by adding TRI Reagent (Merck, T9424) to the fractions at a 1:1 v/v ratio, and then following the manufacturer’s protocol.

### Cell fractionation and isolation of mitoribosomes

Mitochondria and cytosolic fractions were purified as previously described (23) with few modifications. Mitochondrial extracts were loaded onto a 10-30% sucrose gradient prepared as described in the previous section. Mitoribosomes were separated by centrifugation at 38000 rpm for 210 min using a Beckmann SW41 rotor. MAMs isolation was performed as previously described (24).

### Western blot and Immunoprecipitation

Equal amounts of protein from cell lysates were subjected to SDS-PAGE and transferred to a PVDF membrane (Millipore). HeLa GFP and HeLa TRAP1-GFP cells were treated with either 2 μg/mL of harringtonine and 100 μg/mL of emetine for 15 min at 37°C to inhibit cytosolic translation or 30 μM linezolide or 200 ng/mL chloramphenicol for 1 hr at 37°C to inhibit mitochondrial translation. eGFP-fusion proteins were immunoprecipitated with GFP-trap magnetic agarose beads (GFP-trap_MA Chromotek) according to manufacturer’s instructions. The following antibodies were used for WB, immunofluorescence and immunoprecipitation: anti-TRAP1 (Santa Cruz Biotechnology, sc-13557 and Genetex, GTX102017), anti-β-ACTIN (Santa Cruz Biotechnology, sc-69879), anti-puromycin (Merck, MABE343), anti-HSP90 (Santa Cruz Biotechnology, sc-1057), anti-Tubulin (Sigma-Aldrich, T9026), anti-TOM40 (Genetex, GTX133780), anti-TIM44 (Santa Cruz Biotechnology, sc-390755), anti-TIM23 (Santa Cruz Biotechnology, sc-514463), anti-TOM20 (Santa Cruz Biotechnology, sc-17764), anti-GAPDH (Santa Cruz Biotechnology, sc-69778), anti-PHB2 (Santa Cruz Biotechnology, sc-133094), anti-TUFM (Genetex, GTX101763), anti-eEF1a (Millipore, #05-235), anti-IDH2 (Genetex, GTX133078), anti-Rieske (Santa Cruz Biotechnology, sc-271609), anti-STAT1 (Santa Cruz Biotechnology, sc-346), anti-RPS6 (Santa Cruz Biotechnology, sc-74459), anti-RPL19 (Santa Cruz Biotechnology, sc-100830), anti-MRPS5 (Genetex, GTX103930), anti-GFP (Santa Cruz Biotechnology, sc-81045). Total protein normalization has been performed by using No-Stain™ Protein Labeling Reagent (ThermoFisher Scientific, Cat.n. A44717). Images were acquired with a Chemidoc MP imaging system (Bio-Rad), and, where indicated, protein levels were quantified by densitometric analysis using the software ImageJ (25).

### ^35^S Met/^35^S Cys labelling – Puromycilation assay

HeLa Flp-In^™^ T-REx^™^ were seeded in a 6-well plate. HeLa GFP and HeLa TRAP1-GFP cells were induced with 1 μg/mL tetracycline for either 24 hrs (HeLa GFP and HeLa TRAP1-GFP) or 48 hrs (HeLa shGFP and shTRAP1). ^35^S Met/^35^S Cys labeling was performed as follows. Following induction of overexpression or silencing of TRAP1, cells were incubated in cysteine/methionine-free medium (Sigma-Aldrich) for 15 min at 37°C, followed by incubation in cysteine/methionine-free medium containing 50 μCi/mL ^35^S-labeled cysteine/methionine (Perkin-Elmer) for 30 min. Cells were then washed with PBS and lysed. 10 μg of total protein extract was analysed by SDS-PAGE and autoradiography. For the puromycilation assay, following induction, cells were treated with puromycin (1 μg/mL) for 10 min to allow for the puromycin to be incorporated into newly synthesized proteins. Cells were then harvested and total lysates were used for western blot.

### Mitochondrial-specific fluorescent noncanonical amino acid tagging (Mito-FUNCAT)

HeLa shGFP and shTRAP1 cells were seeded in four 15 cm plates. L-Homopropargylglycine (HPG – Thermo Scientific) labeling was performed as follows. Following silencing of TRAP1 for 72 hrs upon induction with 1 μg/mL tetracycline, cells were incubated in cysteine/methionine-free medium (Sigma-Aldrich) for 45 min at 37°C and then treated with 100 μg/mL of cycloheximide and 100 μg/mL of emetine for 15 min at 37°C to inhibit cytosolic translation. Subsequently, 100 μM of HPG was added to the media for 2 hrs at 37°C. After harvesting the cells, mitochondria were isolated by differential centrifugation, as previously described (23). Freshly isolated mitochondria were resuspended in 50 μl of click-it lysis buffer (50 mM TRIS-HCl pH 8, 1% SDS, 250 U/mL Universal Nuclease [Thermo Fisher]) and incubated on ice for 15 min. Mitochondrial extracts were centrifuged at 18’000 rcf for 5 min and protein concentrations were determined by BCA assay (Thermo Fisher). 180 μg of mitochondrial proteins were subjected to a click reaction using a commercial kit (Click-iT Cell Reaction Buffer Kit; Thermo Fisher), with 40 μM Tetramethylrhodamine (TAMRA)-azide (Sigma). According to the manufacturer’s protocol, proteins were purified from the mixture using a MeOH/chloroform approach, after the end of the click reaction. The extracted pellet was dissolved in 20 ul of click-it lysis buffer containing 3% SDS and protein concentrations were determined by BCA assay. Equal amounts of proteins were loaded on a 15% Tris-Glycine gel. Fluorescent signals in the gel were analyzed using a Chemidoc MP imaging system (Bio-Rad). Total protein levels were detected using the No-Stain Protein Labeling Reagent (Thermo Scientific). Click chemistry and immunostaining of fixed cells have been performed as follows. After HPG labeling, coverslips were fixed in 4% paraformaldehyde and pre-permeabilized with a 0.0005% digitonin solution. Then cells were fully permeabilized with 0.1% Triton X-100 for 5 min at room temperature and subsequently incubated with the click reaction buffer for 30 min according to the Click-iT Cell Reaction Buffer Kit (Thermo Fisher Scientific). After washing with 2% BSA in PBS solution, mitochondria were immunostained with anti-TOM20 primary antibody (Santa Cruz Biotechnology, sc-17764) and then cells were subsequently labelled with Alexa Fluor488-conjugated secondary antibody (Invitrogen). The coverslips were mounted using mowiol, and images were acquired using confocal microscope Zeiss LSM700.

### *In vitro* translation assay (including transcription and capping of the GFP mRNA)

The Wheat Germ extract kit used to synthesize eGFP mRNA and the RTS 100 *E. coli* HY Kit used to synthesize EmGFP mRNA were purchased from Promega (Cat. No. L4380) and Biotechrabbit (Cat. No. BR1400102), respectively. For the Wheat Germ translation assay, we assembled each reaction as follows: 5 μL of Wheat Germ Extract, 1 mM of amino acids mixture minus Methionine, 1 mM of amino acids mixture minus Leucine, 8 units of RNasin ribonuclease inhibitor, in a final volume of 10 μL. For the *E. coli* transcription/translation assay, we assembled each reaction as follows: 2.4 μL of *E. coli* lysate, 2 μL of reaction mix, 2.4 μL of amino acids, 0.2 μL of Methionine, 1 μL of reconstitution buffer, in a final volume of 10 μL. mRNAs were added to reactions at a final concentration of 21.95 ng/μL. Where indicated, TRAP1 recombinant protein was added to the reaction at a final concentration of 0.3 μg/μL in Wheat Germ translation assay and 0.2 μg/μL in *E. coli* transcription/translation assay. eGFP and EmGFP fluorescence increases were recorded at 535 nm, at 1-min intervals for 5 hrs at 32° C and at 30-sec intervals for 3 hrs at 30°C, respectively. mRNAs were obtained using the MEGAscript Kit (ThermoFisher Scientific, Cat. No. AM1334) and prepared according to the manufacturer’s manual. Capped eGFP transcript was synthesized by adding a cap analog directly to the MEGAscript reaction.

### RNA extraction and RT-qPCR analysis

Total RNA extraction was performed using the TRI Reagent (Merck, product code T9424) following the manufacturer’s instruction. For first-strand synthesis of cDNA, 1μg of RNA was used in a 20-μl reaction mixture by using a SensiFast cDNA synthesis kit (Bioline). For real-time PCR analysis, 0.4μL of cDNA sample was amplified by using the SensiFast Syber (Bioline) in an CFX Opus 96 Real-Time PCR Instrument (Bio-Rad Laboratories GmbH, Segrate, Italy). The reaction conditions were 95 °C for 5 min followed by 45 cycles of 15 sec at 95 °C and 1 min at 60 °C. The sequence of the oligos used are listed below in 5’-3’ direction:

ACTIN forward: CCTTTGCCGATCCGCCG
ACTIN reverse: AATCCTTCTGACCCATGCCC
ATP5B forward GGACTATTGCTATGGATGGTACAG
ATP5B reverse CCATGAACTCTGGAGCCTC
ATP5MJ forward CTGCGCCAAGATGCTTCAAA
ATP5MJ reverse GGTTAGTGATGACCAGGAGCA
ATP5MK forward GACACCAGCTGCGGAATTTG
ATP5MK reverse ATGCTTCCATATGTGGCCAGT
ATP5PO forward CTCGGGTTTGACCTACAGCC
ATP5PO reverse TGTGGCATAGCGACCTTCAA
ATP6 Forward ACCACAAGGCACACCTACAC
ATP6 Reverse TATTGCTAGGGTGGCGCTTC
ATP8 Forward ACTACCACCTACCTCCCTCAC
ATP8 Reverse GGATTGTGGGGGCAATGAATG
CO1 Forward AATCATCGCTATCCCCACCG
CO1 Reverse CAGAGCACTGCAGCAGATCA
CO2 Forward CCGTCTGAACTATCCTGCCC
CO2 Reverse GAGGGATCGTTGACCTCGTC
CO3 Forward ACCCTCCTACAAGCCTCAGA
CO3 Reverse TGACGTGAAGTCCGTGGAAG
CYB Forward GTCCCACCCTCACACGATTC
CYB Reverse TGGGAGGTGATTCCTAGGGG
LUCIFERASE forward TACAACACCCCAACATCTTCGA
LUCIFERASE reverse GGAAGTTCACCGGCGTCAT
ND1 Forward GCTCTCACCATCGCTCTTCT
ND1 Reverse CCGATCAGGGCGTAGTTTGA
ND2 Forward AGCACCACGACCCTACTACT
ND2 Reverse TGGTGGGGATGATGAGGCTA
ND3 Forward GCCCTCCTTTTACCCCTACC
ND3 Reverse GCCAGACTTAGGGCTAGGATG
ND4 Forward TTCCCCAACCTTTTCCTCCG
ND4 Reverse TGGATAAGTGGCGTTGGCTT
ND4L Forward TCGCTCACACCTCATATCCTC
ND4L Reverse AGGCGGCAAAGACTAGTATGG
ND5 Forward GCTTAGGCGCTATCACCACT
ND5 Reverse TGCAGGAATGCTAGGTGTGG
ND6 Forward GGAGGATCCTATTGGTGCGG
ND6 Reverse CCTATTCCCCCGAGCAATCTC
RPL36A Forward GTGGCTATGGTGGGCAAACTA
RPL36A Reverse ACTGGATCACTTGGCCCTTTCT
12S rRNA Forward AAGCGCAAGTACCCACGTAA
12S rRNA Reverse GGGCCCTGTTCAACTAAGCA

### Förster resonance energy transfer (FRET) assay by fluorescence lifetime imaging (FLIM)

In FRET experiments, the TRAP1-GFP fusion protein or a cy2 conjugated to a secondary antibody were used as donor, while cy3 conjugated to a secondary antibody was used as acceptor. The cells were fixed in 2% paraformaldehyde (PFA), mounted on a slide and analyzed using a TCS SMD FLIM Leica SP5 microscope (Leica, Wetzlar, Germany) equipped with a 63X/1.4 NA objective, to measure FRET efficiency (E_FRET_). E_FRET_ varies as the sixth power of the distance (r) between the two molecules according to the following formula: E_FRET_ = 1/[(1 +r/R_0_)^6^], where R_0_ is the distance corresponding to E_FRET_ = 50%, which can be calculated for any pair of fluorescent molecules. For distances less than R_0_, FRET efficiency is close to maximal, because of the 1/r^6^ dependence, whereas for distances greater than R_0_ the efficiency is close to zero. E_FRET_ by FLIM was calculated with the following formula: E_FRET_: 1-(t_DA_/ t_D_), where t_DA_ is the donor lifetime in presence of the acceptor, while t_D_ is the lifetime of the donor alone.

### Duolink in situ proximity ligation assay

Duolink in situ proximity ligation assay (Navinci - NF.MR.100) was performed according to the manufacturer’s instructions. Briefly, cells were seeded on coverslips, fixed, permeabilized and hybridized with primary antibodies. After one day, cells were hybridized with secondary antibodies conjugated with the PLA probes (PLUS and MINUS), and then subjected to ligation and rolling circle amplification using fluorescently labeled oligonucleotides. Cells were washed and mounted on slides using a mounting media with DAPI to detect nuclei and the signal was detected by confocal microscopy analysis. For proximity ligation assays, the following antibodies were used: anti-TRAP1 (sc-13557 and GTX102017), anti-TUFM (GTX101764), anti-MRPL12 (Genetex, GTX114731), anti-TOM40 (GTX133780); anti-TOM20 (sc-17764); anti-PHB2 (sc-133094); anti-HSP90 (sc-1057); anti-TIM44 (sc-390755); anti-phospho-S6 Ribosomal Protein (Ser240/244) (Cell Signaling Technology #2215). Image acquisition was performed by confocal laser-scanning microscopy using Zeiss 510 LSM from Carl Zeiss Microimaging (Oberkochen, Germany) or by using the Leica Thunder Imaging System (Leica Microsystems) equipped with Leica DFC9000GTC camera and a planapo 63x oil immersion (NA 1.4) objective lens. Fluorescence LED light source and appropriate excitation and emission filters were used. Images were acquired taking Z-slices from the top to the bottom of the cell by using the same setting (LED source power, exposure time) and the Small Volume Computational Clearing (SVCC) mode for the different cell lines and in all experimental conditions.

### Stopped-flow FRET assay

70S Initiation complex labeled with Cy3 on protein L11 (Cy3-70SIC) and E-348C-EF-Tu labeled with the QSY9 fluorescence quencher (QSY-EF-Tu) were prepared as described (26). Ternary complex (TC) was obtained incubating in buffer QSY-EF-Tu, yeast Phe, GTP, EP, PK for 5 min at 37°C. Measurements of Cy3-L11 fluorescence were carried out at stopped-flow fluorometer in the absence of TRAP1 (red trace, positive control), and with TRAP1 pre-incubated with either Cy3-70SIC (blue trace) or QSY-TC (purple trace). A negative control (black trace) was performed using a wild type EF-Tu that is unable to quench Cy3 fluorescence.

### Gene co-expression, Gene Set Enrichment and Network analyses

Gene co-expression analyses, defined as similarity of changes in gene expression patterns between genes of interest, was performed by using COXPRESdb (https://coxpresdb.jp; (27)). Pearson’s correlation coefficient was used as a measure of gene co-expression. Gene set enrichment analyses of the co-expression gene list (Supplementary Table 1) was performed using Enrichr (https://amp.pharm.mssm.edu/Enrichr/; (28)). Network analysis on the 100 genes with the highest degree of co-expression with TRAP1 has been performed by using STRING (29).

## RESULTS

### TRAP1 is associated with both the cytosolic and mitochondrial protein synthesis machineries

We have previously demonstrated that the extra-mitochondrial pool of TRAP1 is loosely associated to the endoplasmic reticulum membrane, facing the cytosol (30), and is bound to ribosomes and translation factors (17). TRAP1 attenuates the rate of protein synthesis thus allowing protein quality control and reducing co-translational ubiquitination and degradation of nascent proteins (14). We further characterized the association of TRAP1 to the translational machinery in HeLa cells by polysome profiling, followed by immunodetection of TRAP1 protein in the resulting fractions. Polysome profiling absorbance, measured at 254 nm, indicates the sedimentation of the particles: fractions 1 and 2 free cytosolic proteins or light complexes; fractions from 3 to 6 ribosomal subunits (60S, 40S) and monomer (80S); fractions from 7 to 12 polysomes. The results of these experiments allowed us to demonstrate that TRAP1 is associated with polysomes: in fact, TRAP1 was found in the polysomal fractions (Fig. 1A), whereas treatment with the translation initiation inhibitor harringtonine, which leads to ribosome run-off, reducing the amount of polysomes accumulating in the monosomal fraction, abolished the signal. This finding confirms that the presence of TRAP1 in the collected fractions is actually linked to the presence of actively translating polysomes (Fig. 1B). Similar results were obtained by disassembling polysomes by treating the cell lysate with EDTA (Supplementary Fig. 1A) or by treating cells with the protein synthesis inhibitor puromycin (Supplementary Fig. 1B). The absence from polysomal fractions of cytosolic proteins that are not expected to bind ribosomes (GAPDH) confirms the quality of the fractionation. We observed contamination of other cellular components in the bottom fraction, similar to those documented by others (31), therefore we excluded the last fraction from subsequent analyses. The predominant mitochondrial localization of TRAP1 protein (Fig. 1C) prompted us to verify whether an interaction of TRAP1 with the mitochondrial translation apparatus would occur, analogously to what demonstrated for the cytoplasmic translational machinery. To this aim, we isolated and fractionated mitochondrial ribosomes (referred to as mitoribosomes hereafter) from HeLa cells, and verified the presence of TRAP1 in the resulting fraction by immunoblot. As a result, TRAP1 was clearly detected in the mitoribosomal fractions along with the mitochondrial ribosomal protein uS5m and the mitochondrial translation elongation factor EF-TuMT, whereas the mitochondrial enzyme isocitrate dehydrogenase 2 (IDH2) is totally absent (Fig. 1D). The molecular chaperone HSP90, highly homologous toTRAP1, also present in cancer cell mitochondria (Fig. 1C), is also visible in some of the mitoribosomal fraction, but not at the same extent as TRAP1 (Fig. 1D). As for the polysome profiling, the last collected fraction at the bottom of the gradient showed contamination of other cellular components, therefore it was excluded from subsequent analyses (see next section). Additionally, we performed a proximity ligation assay (PLA) in HeLa cells using antibodies against TRAP1 and the mitochondrial ribosomal protein bL12m. Results showed that TRAP1 binds to mitochondrial ribosomes in HeLa cells, with an average of 28 spots/cell (Fig. 1E).

**Figure 1:**
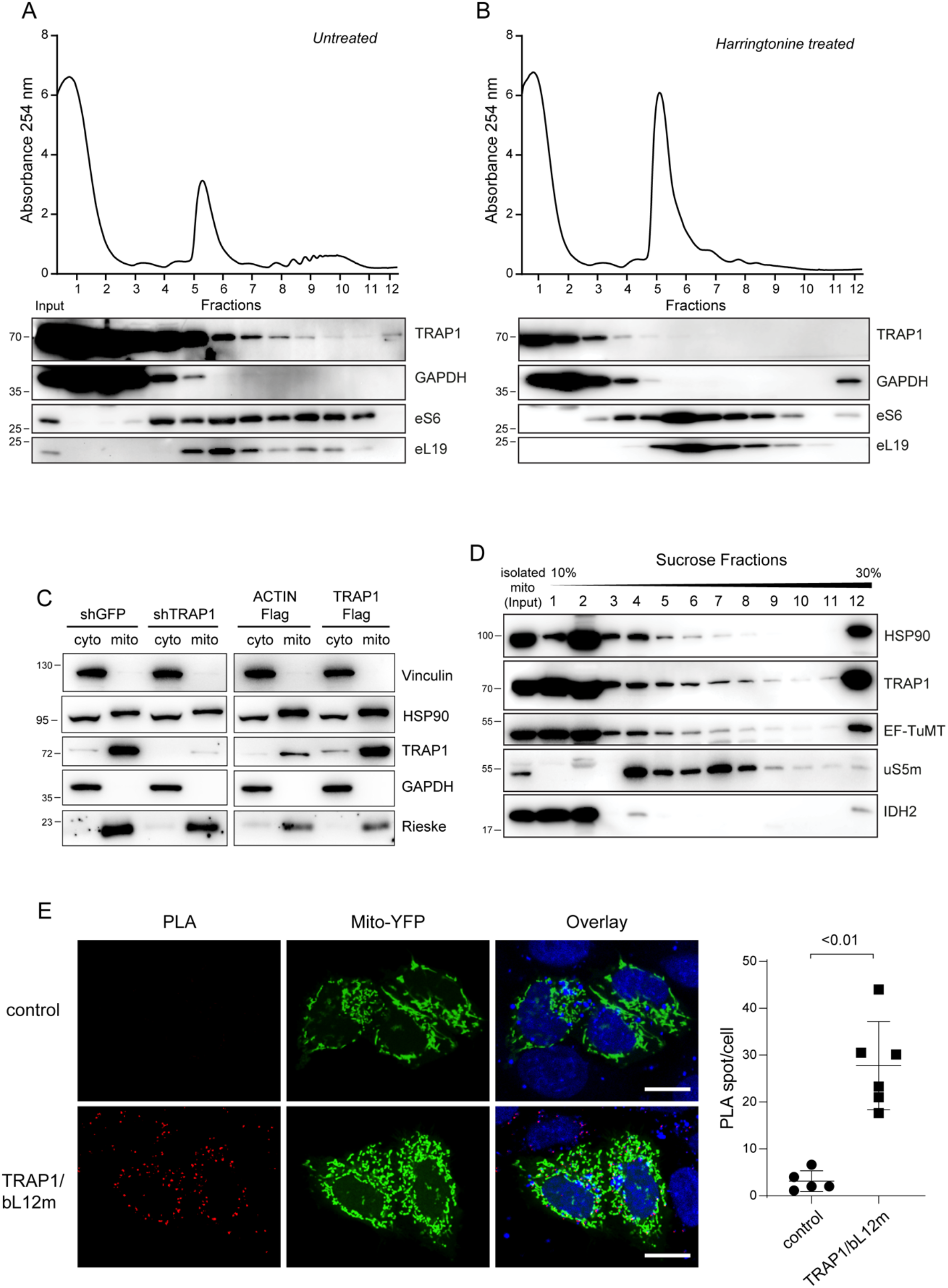
TRAP1 is associated to both cytosolic and mitochondrial ribosomes. (**A-B**) Polysome profiling absorbance, measured at 254 nm, of HeLa cell extracts, from untreated cells (A) or following a 5-minute treatment with 2 μg/mL harringtonine (B). Proteins from each fraction were analysed by WB with the indicated antibodies. (**C**) Subcellular fractionation of HeLa cells showing the presence of indicated proteins into cytosolic (cyto) and mitochondrial (mito) fractions. Vinculin and GAPDH have been used as markers of cytosol and Rieske protein as markers of mitochondria. (**D**) HeLa cell mitochondria were isolated, lysed and loaded onto a 10-30% linear sucrose gradient, followed by fractionation. Proteins were precipitated from the resulting fractions and subjected to western blot with indicated antibodies. (**E**) Representative image of PLA showing the interaction of Hela mitochondria. Positive signals of interaction are shown as red dots, nuclei are stained with DAPI (blue), mitochondria are marked by the mitochondria-directed YFP (green). Scale bar: 10 μm. The graph shows the average number of PLA spot/cell, with a p-value representing the statistical significance based on the Student’s t-test (n=6).

### TRAP1 regulates protein synthesis both in cytosol and mitochondria

To elucidate the molecular mechanisms underpinning TRAP1 contribution to protein synthesis regulation, we characterized its function in a stable, inducible HeLa cell system. First of all, we confirmed in this model that modulation of TRAP1 expression levels, by induction of shRNA-mediated silencing or overexpression of a TRAP1-GFP protein, affected the total cellular protein synthesis rate, as measured by incorporation of ^35^S Met/^35^S Cys. We observed that TRAP1 silencing yielded increased incorporation, whereas TRAP1-GFP overexpression resulted in reduced incorporation compared to unfused GFP-expressing cells (Fig. 2A). Similarly, incorporation of puromycin into proteins is reduced upon TRAP1-GFP overexpression and increased upon shRNA-mediated silencing (Supplementary Fig. 2A).

**Figure 2:**
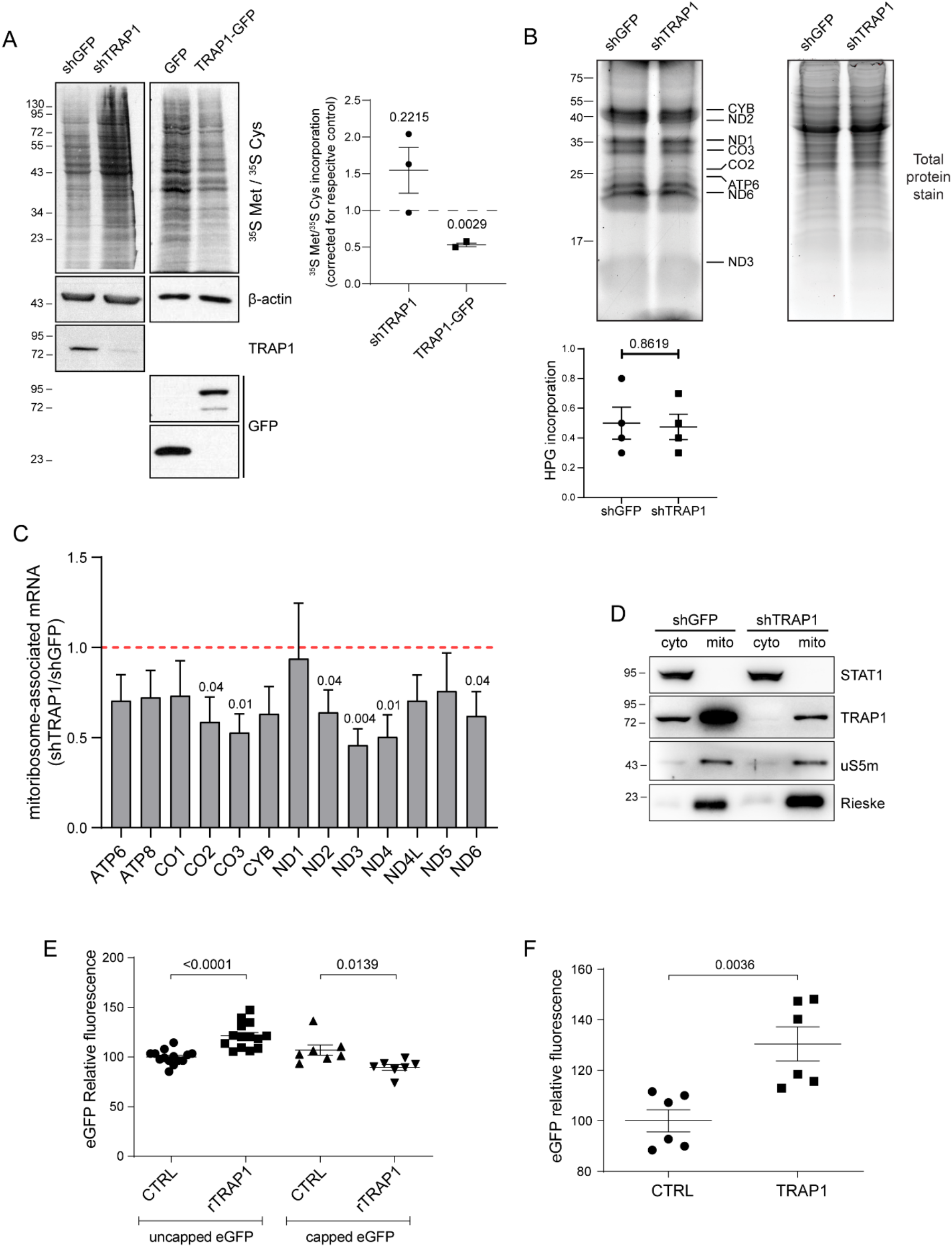
TRAP1 expression has opposite effects on total cell vs. mitochondrial mRNA translation. (**A**) Representative autoradiography of total lysates from cells labelled with ^35^S Met/^35^S Cys, following tetracycline-induced induction of TRAP1-directed shRNA and control shRNA (72 hrs) cells or of TRAP1-GFP and unfused control GFP (24 hrs) cells, with relative densitometric band intensities and analysis (right panel). The p-values in the graph indicate the statistical significance based on the Student’s t-test (n=3). (**B**) Mito-FUNCAT-gel. Expression of GFP (control)-directed and TRAP1-directed shRNAs was induced in HeLa cells with tetracycline 72 hrs before labeling with 100 μM HPG-alkyne for 2 hrs. The resulting lysates were subjected to a click reaction with a TAMRA-azide, loaded for SDS-PAGE and detected at 550 nm. (**C**) RT-qPCR performed on RNAs extracted by mitoribosomal fractions (4–11) isolated from HeLa cells 72 hrs after induction of shGFP/shTRAP1. The amount of mitoribosome-associated mRNA in the two samples has been normalized on 12S rRNA and corrected for its total expression level. Data are expressed as mean **±** SEM (n=5). Numbers above bars represent the statistical significance (p-value) based on the one-sample t-test. (**D**) Subcellular fractionation of HeLa cells showing the presence of indicated proteins into cytosolic (cyto) and mitochondrial (mito) fractions. (**E**) eGFP in vitro translation using wheat germ extracts. eGFP mRNA was added to reactions at a final concentration of 21.95 ng/μL. Where indicated, 0.3 μg/μL of TRAP1 recombinant protein was added to the reaction. Data are expressed as mean **±** SEM (n=14 for the translation of uncapped eGFP mRNA; n=7 for the translation of capped eGFP mRNA). The two-tailed p-value represents the statistical significance based on the Student’s t-test. (**F**) eGFP *in vitro* translation using *E. coli* extracts. eGFP mRNA was added to reactions at a final concentration of 21.95 ng/μL. Where indicated, 0.2 μg/uL of TRAP1 recombinant protein was added to the reaction. Data are expressed as mean **±** SEM (n=6). The p-value represents the statistical significance based on the Student’s t-test.

Since TRAP1 is mainly localized into mitochondria (Fig. 1C and (32)), we further characterized its regulatory role in a mitochondrial translation assay in whole cells by using the Mito-FUNCAT method (33). The assay showed that, in these conditions, there is no difference in the incorporation of the amino acid analog L-Homopropargylglycine (HPG) in the mitochondrial-encoded proteins upon TRAP1 silencing or overexpression (Fig. 2B, Supplementary Fig. 2B-C), on a background of overall equal mRNA abundance (Supplementary Fig. 2D). However, RT-qPCR performed on RNAs extracted by isolated mitochondrial ribosomes in the absence of translation inhibitors (such as cycloheximide, emetine) showed that TRAP1 silencing reduces the association of the mitochondria-encoded transcripts to mitoribosomes (Fig. 2C), suggesting reduced translation. Quantification of mitoribosome-associated mRNAs was corrected on their relative expression levels, and normalized on 12S rRNA, a structural constituent of the ribosome, which rules out a contribution from ribosome biogenesis. This is further confirmed by equal expression levels of the mitochondrial ribosomal protein uS5m in the mitochondria isolated from shGFP and shTRAP1 HeLa cells (Fig. 2D). Interestingly, one key difference with the Mito-FUNCAT approach lays in the crucial requirement of the Mito-FUNCAT method to inhibit cytosolic translation with cycloheximide/emetine, which we believe hampers the possibility to detect differences in mitochondrial translation upon TRAP1 knockdown/overexpression, given its dual role in the two compartments and in the crosstalk between the two, as further explored in the next section. This observation is in line with previous studies performed in yeast (2), demonstrating a unidirectional crosstalk between cytosolic and mitochondrial translation.

To further characterize TRAP1 effects on translation, and taking advantage of our previous data on TRAP1 regulation of capped/uncapped mRNA translation (32), the mechanistic contribution of TRAP1 to mRNA translation was evaluated in an *in vitro* system upon the addition of 5’ methylguanosine cap. As shown in Figure 2E, the addition of a recombinant mature TRAP1 protein in wheat germ extracts reduces the translation of an *in vitro*-transcribed GFP mRNA when a 5’ methylguanosine cap is added to the transcript, whereas it increases translation of the uncapped mRNA, thus confirming not only TRAP1 direct role in translation, but also that high expression of TRAP1 correlates with a higher IRES/cap-dependent translation ratio, in agreement with our previous observations (20). Accordingly, given the similarities between mitochondrial and prokaryotic translation, we found that TRAP1 addition increases GFP reporter protein synthesis in *E. coli* extracts (Fig. 2F), showing that TRAP1 actually increases the translation of leaderless mRNAs. These findings also support the results indicating a reduced association of mitochondrial transcripts to mitoribosomes upon TRAP1 silencing (Fig. 2C).

### TRAP1 couples cytosolic with mitochondrial translation

The mitochondrial respiratory complexes are composed by both nuclear-encoded and mitochondrial-encoded proteins. The former are synthesized into the cytosol and then have to be imported into the mitochondrion, while the latter are synthesized within the organelle through a mechanism largely dependent on the assembly of the complex, and therefore on the availability of the nuclear-encoded components (34). Of note, additional to TRAP1 involvement to both the translation processes, previous results from a quantitative proteomic analysis (35) showed the mitochondrial protein import channel component TOM40 as the second most significant interactor among the TRAP1 protein partners in HeLa cells. Here, to provide further evidence for this interaction, two additional approaches were used: 1) proximity ligation assay showing that endogenous TRAP1 and TOM40 produce an average of 13 spots/cell (Fig. 3A); 2) co-immunoprecipitation experiments showing that a binding between the two proteins is present when the TRAP1-GFP protein is isolated from inducible HeLa cells by a GFP-trap employing highly specific nanobodies, whereas no signal is observed with other components of the TIM-TOM complex (TOM20, TIM23, TIM44), or mitochondrial proteins (PHB2) thus supporting the specificity of this binding (Fig. 3B). Similar results were obtained by PLA (Supplementary Fig. 3A). However, TRAP1 also interacts with mitochondrial protein localized into the matrix, which, according to mass spectrometry data, includes the mitochondrial translation elongation factor EF-TuMT (36,37). This latter interaction was confirmed by proximity ligation assay in HCT116 cells (Fig. 3C) and HeLa cells (Supplementary Fig. 3B). The highly homologous HSP90 does not show proximity ligation with EF-TuMT (Supplementary Fig. 3B), further supporting the specificity of TRAP1-EF-TuMT binding. The binding between TRAP1 with cytosolic translation factors (Supplementary Fig. 3C and (17)) and ribosomes, protein import channel components and mitochondrial translation factors and ribosomes prompted us to evaluate whether these interactions are dependent on active protein synthesis. To this aim, we treated cells with cytosolic translation initiation (harringtonine) or elongation (emetine) inhibitors, or with inhibitors of mitochondrial translation initiation (linezolid) or elongation (chloramphenicol). Co-immunoprecipitation performed in these conditions strikingly showed that inhibition of cytosolic protein synthesis dramatically reduces the binding between TRAP1 and TOM40, while TRAP1-eEF1A and TRAP1-EF-TuMT interactions increased (Fig. 3D). Conversely, inhibition of mitochondrial translation yielded no effect on the binding of either cytosolic or mitochondrial partners (Fig. 3E). Again, and in agreement with the observation in yeast (2), this result confirms the unidirectional nature of the process: i.e. cytosolic translation transduces a signal to mitochondrial translation apparatus, while mitochondrial translation is unable to reverse a signal to the cytosolic ribosomes, and TRAP1 takes part to this mechanism.

**Figure 3:**
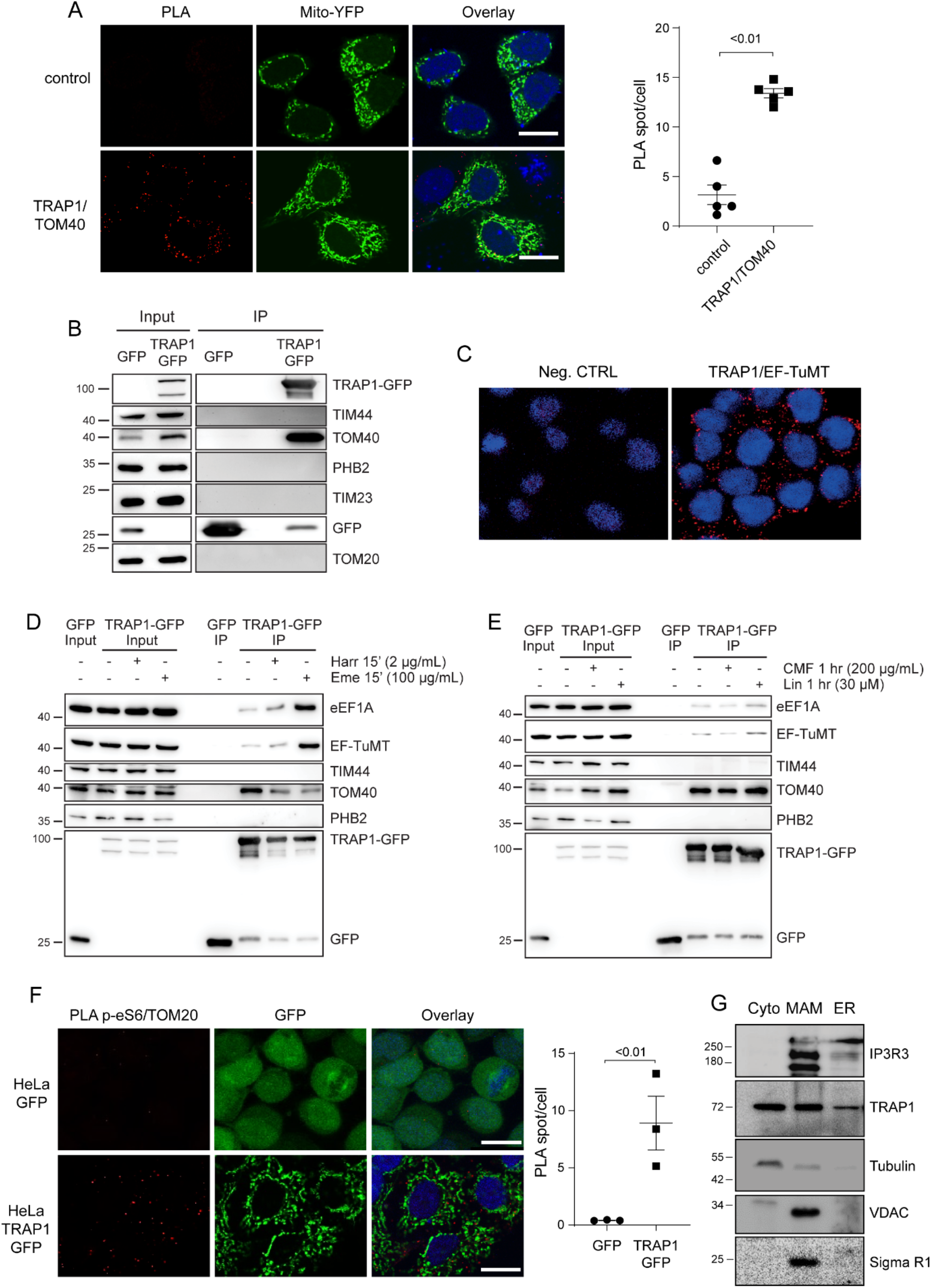
TRAP1 associates with the mitochondrial protein import machinery and favors localized translation. (**A**) Representative images of PLA positivity between TRAP1 and TOM40. Positive signals of interaction are shown as red dots, nuclei are stained with DAPI (blue), mitochondria are marked by the mitochondria-directed YFP (green). Scale bar: 10 μm. The graph shows the average number of PLA spots/cell, with a two-tailed p-value representing the statistical significance based on the Student’s t-test (n=5). (**B**) Immunoprecipitation of unfused GFP and TRAP1-GFP performed in HeLa cells following 24 hrs induction of GFP and TRAP1-GFP. Total lysates were incubated with GFP-trap beads to isolate the proteins and the resulting samples were immunoblotted with indicated antibodies. (**C**) Representative image of PLA showing the interaction of TRAP1 with EF-TuMT in HCT116 cells. Positive signals of interaction are shown as red dots, nuclei are stained with DAPI (blue). Negative control has been obtained by hybridizing cells with TRAP1 antibody only. (**D-E**) Immunoprecipitation of unfused GFP and TRAP1-GFP performed in HeLa cells following 24 hrs induction of GFP and TRAP1-GFP. Where indicated, cells were treated for 15 min with emetine (100 μg /mL) or harringtonine (2 μg/mL), or for 1 hr with chloramphenicol (200 μg /mL) or Linezolid (30 μM). Total lysates were incubated with GFP-trap beads to isolate the proteins and the resulting samples were immunoblotted with indicated antibodies. (**F**) Representative image of PLA showing the interaction of TOM20 with phosphorylated (active) ribosomal protein eS6 in HeLa cells following 24hrs induction of TRAP1-GFP or unfused GFP. Positive signals of interaction are shown as red dots, nuclei are stained with DAPI (blue). Scale bar: 10 μm. The graph shows the average number of PLA spots/cell, with a p-value representing the statistical significance based on the Student’s t-test (n=3). (**G**) Cytosolic, MAM, ER, fractions isolated from HCT116 cells probed with the indicated antibodies.

TOM20, another mitochondrial import channel component, is involved in the translation-dependent enrichment of mRNAs encoding mitochondrial proteins near mitochondria, facilitating protein transport into the organelle (4). To evaluate whether TRAP1 might be involved in both the import of mitochondrial proteins and localized protein synthesis, we used a proximity ligation assay: results show that TRAP1-GFP induction in HeLa cells increases the number of proximity ligation spots between TOM20 and actively translating ribosomes, as detected by phosphorylation of the ribosomal protein eS6 (Fig. 3F). In contrast, TRAP1 interference by shRNAs decreased such proximity (Supplementary Fig. 3D). Remarkably, we found that TRAP1 is indeed localized in the so-called mitochondria-associated membranes (MAMs), regions common to all cells in which the endoplasmic reticulum and mitochondria are physically connected (Fig. 3G). These results not only support our previous data on TRAP1 role in protein synthesis control at cytosol/mitochondria interface/crossroad but, for the first time, identify TRAP1 in the MAMs compartment. Altogether, as MAMs are ‘‘hot spots’’ for the intracellular signaling of important pathways (38), the finding of TRAP1 physically associated to these structures provides further support for our previously described functions of TRAP1 in lipid biosynthesis (18), calcium homeostasis (39), reactive oxygen species generation (40), and protein sorting (41).

### TRAP1 regulates translation elongation

To shed light in the mechanisms leading to TRAP1 translational control, we questioned whether the binding to translation elongation factors could explain TRAP1 role in the modulation of protein synthesis. This hypothesis was also supported by the evidence that the modulation of translation elongation rate reduces co-translational protein degradation (42), and this would be consistent with TRAP1 function in preventing ubiquitination of newly synthesized proteins (17,30). Additionally, recent studies performed in yeast show that translation elongation stall increases mRNA localization to mitochondria (43).To this aim, we first investigated TRAP1 binding to the translation elongation factor eEF1G by Fluorescence Lifetime Imaging (FLIM) followed by confocal microscopy analysis, which allows verification of direct protein-protein interactions with high accuracy (44). Our results showed that TRAP1 binds eEF1G in HeLa cells directly, thus also confirming the presence of TRAP1 in the cytosolic compartment and its association with the cytosolic translational machinery (Fig. 4A). Intriguingly, and in agreement with its dual translational control, besides the binding to cytosolic translation elongation factors, TRAP1 also binds the mitochondrial translation elongation factor EF-TuMT, as suggested by mass spectrometry data (36,37), and shown above by proximity ligation assay (Fig. 3C, Supplementary Fig. 3B) and immunoprecipitation (Fig. 3D-E). Therefore, we explored this issue deeply and found that the two proteins are actually bound directly with high efficiency, as confirmed by FLIM in HeLa TRAP1-GFP cells (Fig. 4B). In a related experiment employing a stopped-flow fluorescence assay (21), we demonstrated a direct interaction of TRAP1 with EF-Tu, the bacterial homolog of the mitochondrial EF-Tu (EF-TuMT). This experiment monitors the interaction of a ternary complex, aa-tRNA.EF-Tu-GTP, containing EF-Tu labeled with a fluorescence quencher, with a fluorescent-labeled ribosomal 70S initiation complex containing fMet-tRNA^fMet^ in the ribosomal P-site, as part of the first elongation step which results in formation of dipeptidyl-tRNA. In the positive control, entry of the quencher labeled ternary complex into the ribosomal A-site results in a rapid decrease in fluorescence, which was rapidly recovered upon the EF-Tu.GDP release from the ribosome (Fig. 4C). Addition of TRAP1 inhibits such release, demonstrating the TRAP1:EF-Tu interaction. Inhibition of EF-Tu release, or, more in general, EF1 release, may be a more relevant factor for regulating the rate of cytoplasmic rather than mitochondrial protein synthesis (see Discussion). To evaluate the relevance of these findings in human cells, we set up a SunRiSE assay (45) to assess whether such a mechanism reflects changes in the elongation rate. Briefly, following inhibition of translation initiation with harringtonine, we measured puromycin incorporation into nascent peptide chains at different time intervals. Interestingly, the decay of puromycin labeling following harringtonine treatment was found somewhat steeper in TRAP1 knockdown cells as compared to the shGFP controls (Fig. 4D), indicating a higher speed of run-off of ribosomes from mRNAs, consistent with a higher processing speed of the ribosomes along mRNAs in the absence of TRAP1.

**Figure 4:**
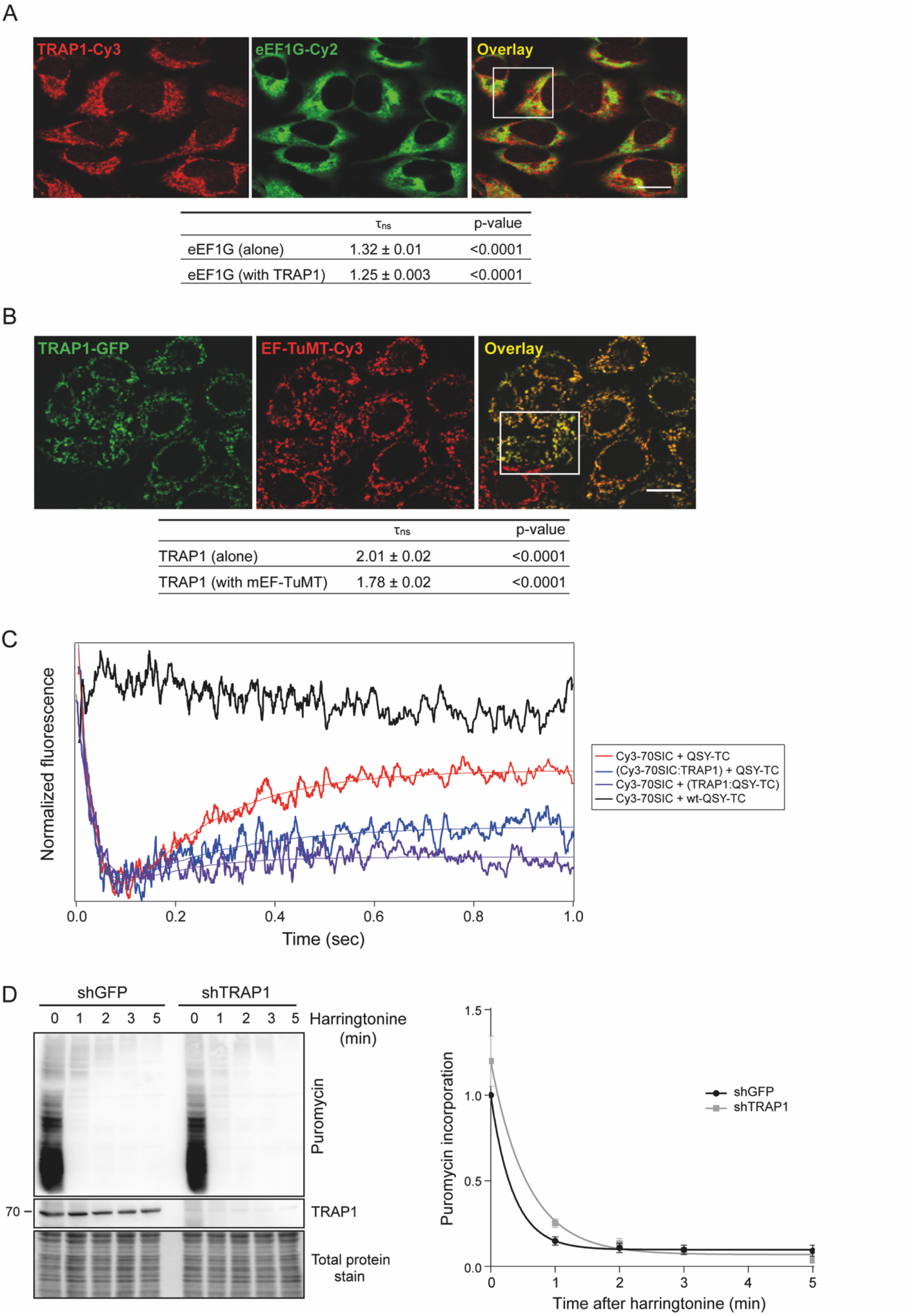
TRAP1 binds both cytosolic and mitochondrial translation elongation factors and slows down elongation rate. (**A**) Fluorescent confocal microscopy analysis of TRAP1-cy3 (acceptor) and eEF1G-cy2 (donor) in HeLa cells. Dipole-dipole energy transfer from the fluorescent donor to the fluorescent acceptor allowed calculating FRET efficiency (E_FRET_ %) as described in Materials and Methods section. The overlay images show the InSet area in which FRET has been analyzed. Scale bar: 10 μm. **τ_ns_** values are expressed as mean **±** SEM. The two-tailed p-value represents the statistical significance based on the Student’s t-test. (**B**) Fluorescent confocal microscopy analysis of TRAP1-GFP (donor) and EF-TuMT-Cy3 (acceptor) in TRAP1-GFP inducible HeLa cells. Dipole-dipole energy transfer from the fluorescent donor to the fluorescent acceptor allowed calculating FRET efficiency (E_FRET_ %) as described in Materials and Methods section. The overlay images show the InSet area in which FRET has been analyzed. Scale bar: 10 μm. **τ_ns_** values are expressed (**C**) TRAP1 inhibits EF-Tu release from 70S initiation complex (70SIC) in stopped-flow assays. TRAP1 recombinant protein was preincubated with a Ternary Complex (TC) in which EF-Tu is labeled with the QSY9 fluorescence quencher (QSY-TC, purple trace) and the resulting solution was rapidly mixed with a Cy3-labeled 70S Initiation Complex (Cy3-70SIC, blue trace). The change in Cy3 fluorescence was monitored using a stopped-flow fluorometer. Upon entering in the A site, the quencher-labeled EF-Tu decreases the Cy3-labeled ribosome fluorescence, whereas its dissociation from the ribosome allows Cy3 fluorescence recovery. Black trace, negative control; red trace, positive control. (**D**) 72 hrs after tet-induction shGFP-directed (control) or TRAP1-directed shRNAs, HeLa cells were treated with harringtonine (2 μg/mL) for the indicated times (0, 1, 2, 3 and 5 min) and subsequently treated with puromycin (10 μg/mL) for 10 min. Cells were lysed and subjected to immunoblotting with anti-puromycin antibody. The graph shows densitometric intensity of the puromycin labeling, normalized on the total protein content (quantified by no-stain labeling, see Methods). Data are represented as mean **±** SEM from 6 independent experiments, with trend lines showing exponential one-phase decay analysis.

Accordingly, a 2-minute treatment with Harringtonine caused a lower decrease in polysomes and increase in monosomes in TRAP1-Flag overexpressing cells compared to the control ACTIN-Flag cells (Fig. 5A), while shTRAP1 cells displayed faster run-off compared to shGFP controls (Fig. 5B). Again, these findings are consistent with the hypothesis that TRAP1 regulates translation at the elongation step. Considering the dual role in the regulation of cytosolic and mitochondrial protein synthesis, we measured the translation speed of mRNA transcripts encoding several mitochondrial ATPase proteins, synthesized by either mitochondrial (ATP6 and ATP8) or cytoplasmic (ATP5 subunits) ribosomes: to this aim, we set up a run-off experiment and determined the ratio between polysome-associated and monosome-associated mRNAs following harringtonine treatment: control (shGFP) and shTRAP1 HeLa cells were treated for 2 minutes with harringtonine and fractionated, and the amount of transcripts associated with polysomal and monosomal fractions of both treated and untreated cells was estimated by qPCR. Figure 5C shows the polysome/monosome ratio for each transcript upon harringtonine treatment, compared to the untreated samples. For the ATP5-encoding transcripts, this ratio is much more reduced in the shTRAP1 cells than in the shGFP controls. In contrast, no significant reduction occurs when mitochondrial-encoded transcripts are analyzed, as expected, since harringtonine does not inhibit mitochondrial ribosomes. Transcripts encoding cytosolic proteins like the ribosomal protein eL42 (gene name RPL36A) respond to harringtonine by reducing their polysome/monosome ratio, but do not display differences in this reduction between shGFP and shTRAP1 cells.

**Figure 5:**
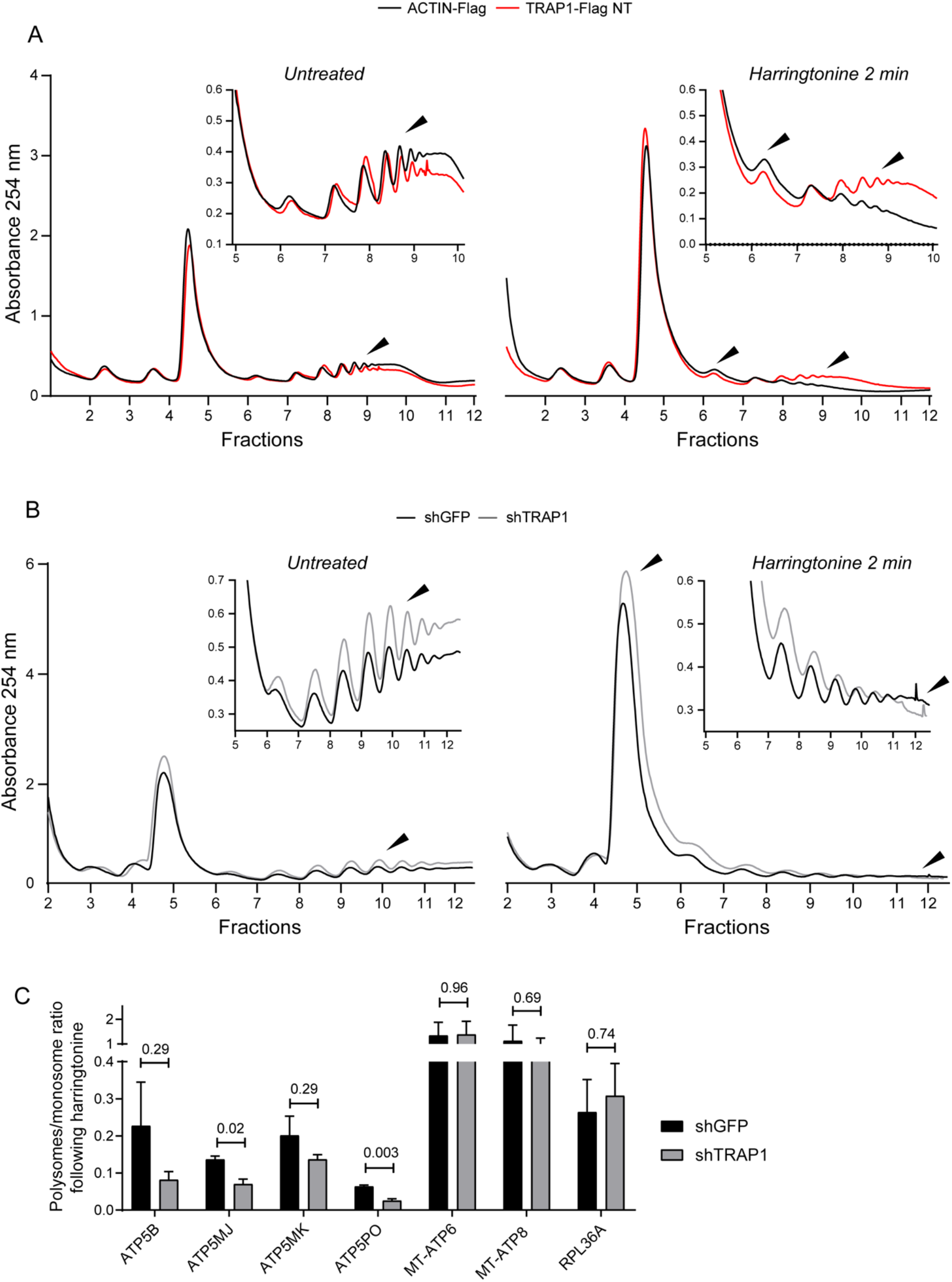
TRAP1 slows down translation elongation by cytoplasmic ribosomes. (**A-B**) Polysome profiling absorbance, measured at 254 nm, of extracts from ACTIN-Flag (control) and TRAP1-Flag overexpressing HeLa cells (A) and from shGFP (control) and shTRAP1 HeLa cells, 24 hrs (A) or 72 hrs (B) after induction, in the absence (left) or presence (right) of 2 μg/mL of harringtonine (cells were treated for 2 minutes and then blocked with cycloheximide). Inset areas show magnification of regions of interest. (**C**) Ratio between polysome-associated and monosome-associated mRNAs following harringtonine treatment (2 μg/mL, 2 min), normalized on the respective untreated samples. The amount of the associated transcripts was measured by RT-qPCR performed on RNAs extracted from pooled monosomal and polysomal fractions, both corrected for an external reference spike-in RNA (luciferase). Data are represented as mean **±** SEM from 3 independent experiments. Number above bars represent the statistical significance (p-value) calculated by a multiple t test.

### TRAP1 is a putative chaperone linked to protein synthesis (CLIPS)

Previous data identified and functionally characterized a selective group of chaperones linked to protein synthesis (CLIPS), co-expressed with components of the translational machinery in yeast (46). Our findings suggest that TRAP1, due to its role in protein synthesis regulation, might behave as the first CLIPS (or CLIPS-like) identified in higher organisms. To test this unprecedently hypothesized role of TRAP1 as a mammalian CLIPS, we performed a co-expression analysis using the software COXPRESdb (27), and found that TRAP1 is significantly co-expressed with several components of the mitochondrial translation apparatus, with 10% of the most 100 coexpressed genes encoding mitochondrial ribosomal proteins (Supplementary Table 1). This analysis builds up discrete clusters of proteins related to mitochondrial translation and metabolism connected by EF-TuMT, mostly constituted by mitochondrial ribosomal proteins, electron transport chain components and assembly factors, and organelle biogenesis and metabolism (Fig. 6A). A gene ontology analysis of biological processes performed through Enrichr (23) on the list of gene co-expressions showed that all top five enriched pathways are correlated with ribosomal RNA and tRNA processing and mitochondrial translation (Fig. 6B). For instance, TUFM, the gene encoding EF-TuMT, is the second most correlated gene (Fig. 6C), and, remarkably, the correlation between TRAP1 and EF-TuMT is conserved when protein levels are compared (Fig. 6D). Consistently, Cluh, an RNA-binding protein involved in the proper cytoplasmic distribution of mitochondria, is one of most TRAP1-coexpressed genes. Cluh specifically binds mRNAs of nuclear-encoded mitochondrial proteins in the cytoplasm and regulates the transport or translation of these transcripts close to mitochondria, playing a role in mitochondrial biogenesis (47).

**Figure 6:**
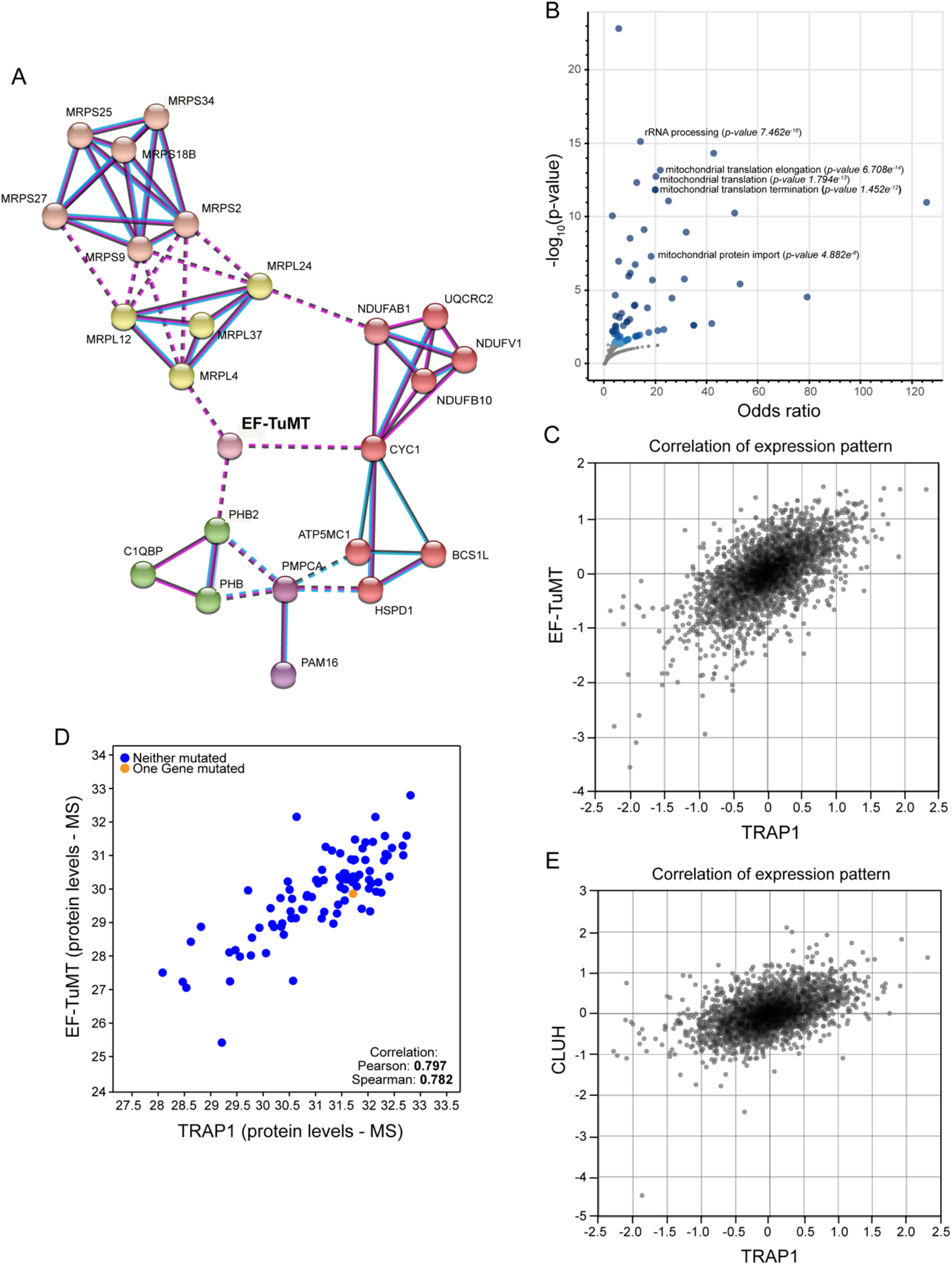
TRAP1 is coexpressed with the mitochondrial translational machinery. (**A**) Network analysis of the top 100 TRAP1-coexpressed genes, generated by STRING using a Markov Cluster (MCL) algorithm. Network nodes represent proteins; edges represent protein-protein associations, by coexpression (black line), experimentally-determined association (pink line), or database-curated association (blue line). Edges between clusters are represented by dotted lines. The red cluster is enriched in mitochondrial electron transport components; pink and yellow clusters are constituted by structural component of the mitochondrial ribosome (small and large subunit, respectively); the green cluster contains the mitochondrial prohibitin complex; the purple cluster is constituted by the association between the mitochondria protein import inner membrane translocase subunit PAM16 and the mitochondrial-processing peptidase subunit alpha PMPCA. (**B**) Gene set enrichment analysis on the list of genes significantly coexpressed with TRAP1 in human tissues. (**C**) Co-expression analysis performed with COXPRESdb between TRAP1 and EF-TuMT (gene name: TUFM). (**D**) Co-expression analysis between TRAP1 and EF-TuMT at protein level according to the Provisional database. (**E**) Coexpression analysis performed with COXPRESdb between TRAP1 and Cluh.

## DISCUSSION

TRAP1, the main mitochondrial member of HSP90 protein family, interacts in mitochondria with respiratory complexes and contributes to the regulation of cellular respiration in cancer cells (7). However, we and others demonstrated that this protein is also partially localized on the outer side of the endoplasmic reticulum, where it is involved in the regulation of protein synthesis through the binding to components of the translational machinery ((17) and this article). Interest in TRAP1 has considerably grown in the past decades due to its contextual effects in different tumor types: it is highly expressed in several cancers and correlated with drug resistance, but is downregulated in specific tumors with predominant oxidative metabolism (48). The interplay between protein synthesis and mitochondrial respiration has been recently investigated (2), but the involvement of this phenomenon in humans is almost unexplored. mRNA localization and localized translation are emerging as important regulatory layers for the expression of mitochondrial metabolism components, which are coordinated by retrograde signaling from the organelle (6). In order to allow coordination of oxidative phosphorylation complexes assembly and activity, mitochondrial and cytosolic translation must be rapidly, dynamically and synchronously regulated (2); this represents a unique challenge for the cells, because their subunits are encoded by both the nuclear and the mitochondrial genomes. In this study, we explored the hypothesis that TRAP1 may function as a “moonlighting” protein, regulating mRNA translation in both the cytosol and mitochondria, upstream to the previously demonstrated modulation of activity respiratory complex II (11), III (13) and IV (14). This favors coordination between the synthesis of mitochondria-destined proteins in the cytosol, their import into the organelle and the synthesis of mitochondrial-encoded proteins in the matrix. This last synthesis is restricted to the translation of only 13 transcripts, exclusively producing components of the electron transport chain. Indeed, we provide the first clear evidence that TRAP1 specifically binds the mitochondrial translation apparatus, facilitating translation of mtDNA-encoded proteins. However, TRAP1 not only participates in both mitochondrial and cytosolic protein synthesis, but is also involved in protein import into mitochondria: TRAP1 specifically binds TOM40 in a translation-dependent manner, and the resulting signal is transduced to the organelle, where TRAP1-EF-TuMT binding is also affected by the activity of cytosolic translation. Noteworthy, we demonstrated for the first time that TRAP1 is localized in the contact sites between the endoplasmic reticulum and mitochondria (the MAM compartment, (38)), not only supporting TRAP1 role in protein synthesis control at cytosol/mitochondria interface but also providing a further support of our previously described functions of TRAP1 in lipid biosynthesis (18), calcium homeostasis (39), reactive oxygen species generation (40), and protein sorting (41). Interestingly, we found that TRAP1 expression is directly correlated to the amount of active ribosomes in the proximity of channel component TOM20. TOM20 is known to mediate the localization of mRNAs encoding mitochondrial proteins to these organelles in a translation-dependent manner (4). Of note, we previously demonstrated that TRAP1 expression is crucial as well for the correct localization of the prion protein Shadoo at the interface between endoplasmic reticulum and mitochondria in neuronal cells (41).

Here for the first time, we present clear evidence that TRAP1 behaves as a translation regulator via direct binding to translation elongation factors. Although translation control has been traditionally attributed mostly to the regulation of the initiation step, evidence is accumulating on the relevance of elongation rate control in the homeostasis of the translation process and its coordination with related pathways (31). Relevant in the light of our findings, it has been recently shown that translation rate is relevant for localization of mRNAs coding for mitochondrial proteins to this organelle, and that translation elongation stall impacts such localization (43). The correlation between translation elongation rate and protein yield is not trivial. Since eukaryotic mRNAs are circularized, potentially allowing terminating ribosomes to preferentially reinitiate on the same transcript, increased elongation rates can result in increased protein yield without changes in ribosome density on a specific mRNA (49). While in a linear model of translation, protein abundance is mainly determined by the initiation rate, in a closed-loop model the continuous re-initiation on the same transcript results in protein yield being significantly determined by the elongation rate. This model fits remarkably well with our data on protein synthesis regulation by TRAP1, showing that it binds a translation elongation factor, inhibits its release from the ribosome and reduces incorporation of amino acids into proteins over the time, without changing the global ribosome loading and therefore the total amount of active polysomes in the cell. Indeed, the presence of TRAP1 only seems to reduce incorporation of amino acids during the translation of capped transcripts, that can be easily circularized, which confirms our model. Of note, although mRNA circularization can occur in mammalian mitochondria, it has been recently shown that it does not appear to play a role in making translatable mRNAs (50). Since uncapped mRNAs may or may not be subject to circularization (51), the model also fits with the evidence indicating that uncapped transcripts are instead more efficiently translated in the presence of TRAP1, since in this case protein yield would be mainly determined by the initiation rate. Additionally, in other cell models, we have already demonstrated that TRAP1-expressing cells show increased levels of eIF2α phosphorylation (which represses only cap-dependent initiation of translation (52)) and a higher ratio of IRES-vs cap-dependent translation (20). The latter mechanism is relevant in cancer development since, among 70 experimentally verified cellular IRES elements (53) (35), a large number are found in cancer-related genes (54). Slowdown of elongation phase also contributes to co-translation protein quality control, contributing to reduce co-translational ubiquitination and degradation of nascent peptides and ensuring stability; in fact, sometimes “less is more” in protein synthesis (55). In this regard, it is noteworthy that we have previously shown how the binding between TRAP1 and the proteasome regulatory subunit component TBP7 in the cytosol reduces global co-translational ubiquitination (19), which critically tunes the expression level of the mitochondria-destined proteins ATP5B and Sorcin isoform B (17); however, the proteasome is absent from mitochondria and, to the best of our knowledge, similar ribosome-bound quality control mechanisms triggering degradation of nascent peptides synthesized into the mitochondria have never been described.

Interestingly, we show that a strong correlation exists between TRAP1 levels and the concomitant expression of the mitochondrial protein synthesis apparatus in human tissues, and, more in general, of proteins involved in mitochondrial biogenesis and metabolism. TRAP1 is also co-expressed with a subset of protein partners, including the translation factors eIF2α and eEF1A, which identifies a cohort of metastatic colorectal carcinomas with a significantly shorter overall survival (56).

Taken together, the present results demonstrate an unprecedented level of complexity in the regulation of cancer cell metabolism, in which both mitochondrial and cytosolic protein synthesis are co-regulated and coordinated with energy metabolism through the contribution of a common molecular chaperone, TRAP1. As cell metabolism is a complex network of interdependent pathways and the correlations between protein synthesis and mitochondrial respiration in human diseases are mostly unknown, the highlighting of bioenergetic pathways, with an eye towards protein synthesis, may provide a novel approach for therapeutic development, adding a further square in the complex puzzle of TRAP1 chaperone functions in cancer cells.

## SUPPLEMENTARY DATA

Supplementary Data are available online.

## ACKNOWLEDGEMENT

We acknowledge prof. John J Rossi for the pFRT-U6tetO plasmid. We acknowledge Elena Dobrikova and Matthias Gromeier (at Duke University Medical Center) for the establishment of the HeLa Flp-In T-Rex cell line. The eGFP alone cloned into pcDNA5/FRT/TO (Invitrogen) was kindly provided by Prof. Matthias Hentze, EMBL/ Heidelberg Univ. ‘Molecular Medicine Partnership Unit’. The inducible Actin-Flag HeLa cell line was kindly provided by Prof. Alfredo Castello, University of Glasgow. We acknowledge microscopy facility at the Department of Molecular Medicine and Medical Biotechnology – University of Napoli “Federico II” for the support in microscopy analyses.

---

**Supplementary Figure 1:**
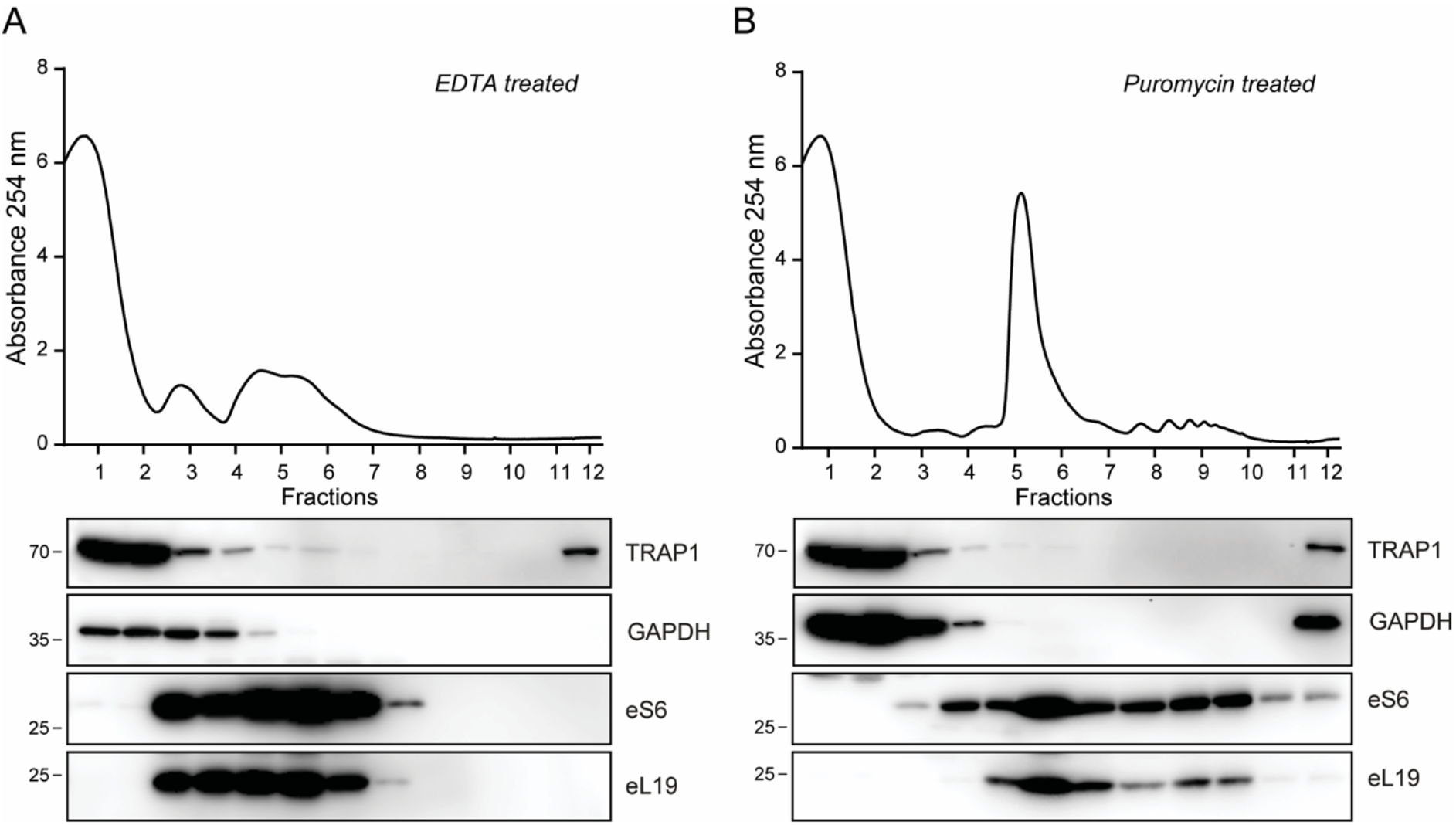
(**A-B**) Polysome profiling absorbance, measured at 254 nm, of HeLa cell extracts, in the presence or absence of 30 mM EDTA (A) or following a 15-minute treatment with 100 μg/mL puromycin (B). Proteins from each fraction were analysed by WB with the indicated antibodies.

**Supplementary Figure 2:**
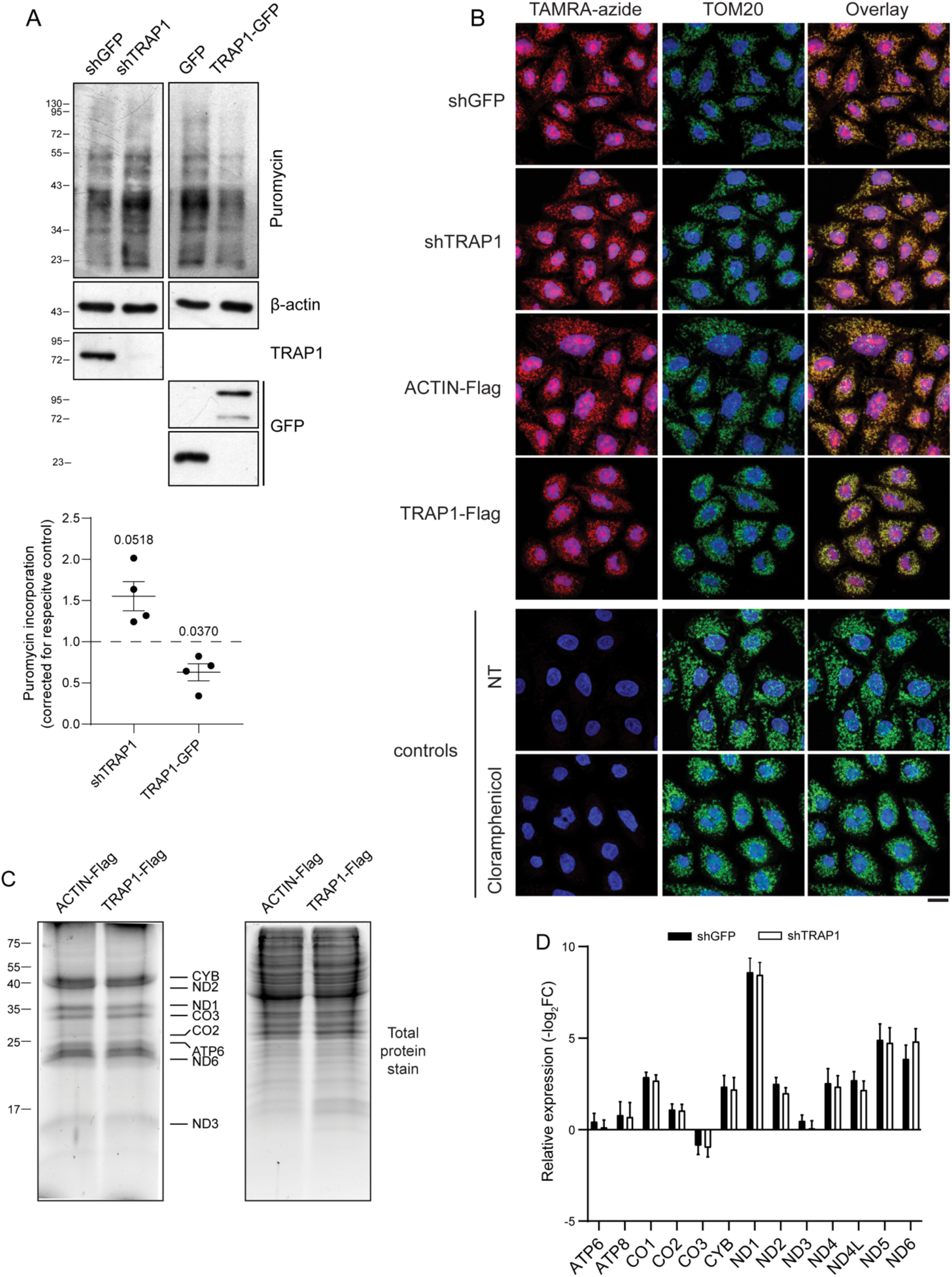
(**A**) Following tetracycline-induced induction of TRAP1-directed shRNA and nontargeting control shRNA (72 hrs) or of TRAP1-GFP and unfused control GFP (24 hrs), HeLa cells were treated with puromycin (1 μg/mL, 15 min). Representative immunoblots of total cellular lysates with indicated antibodies are shown (left panel), with relative densitometric band intensities and analysis (right panel). The p-value in the graph indicate the statistical significance based the Student’s t-test (n=4). (**B**) Mitochondrial translation products were labeled using HPG in both shTRAP1 and TRAP1-Flag HeLa cells and in control cells (shGFP and Actin-Flag, respectively). TOM20 immunostaining was used to confirm the mitochondrial localization of the HPG signal visualized by a copper-catalyzed cycloaddition reaction (click) to TAMRA-azide. We used cells that were not incubated with HPG (NT) or those treated with chloramphenicol as negative controls. Scale bar 10 μm. (**C**) MITOFUNCAT-gel. Expression of Actin-Flag and TRAP1-Flag was induced in HeLa cells with Tetracycline 24 hours before labeling with 100 μM HPG-alkyne for 2 hours. The resulting lysates were subjected to a click reaction with a TAMRA-azide, loaded for SDS-PAGE and detected at 550 nm. (**D**) RT-qPCR of the 13 mitochondrial-encoded protein coding-RNAs upon induction (72 hours) of GFP-directed (control) and TRAP1-directed shRNAs in HeLa cells. Total RNAs were extracted from isolated mitochondria. Data are expressed as mean ± S.E.M. from four independent experiments with technical triplicate each. The 12S rRNA was used as the internal control.

**Supplementary Figure 3:**
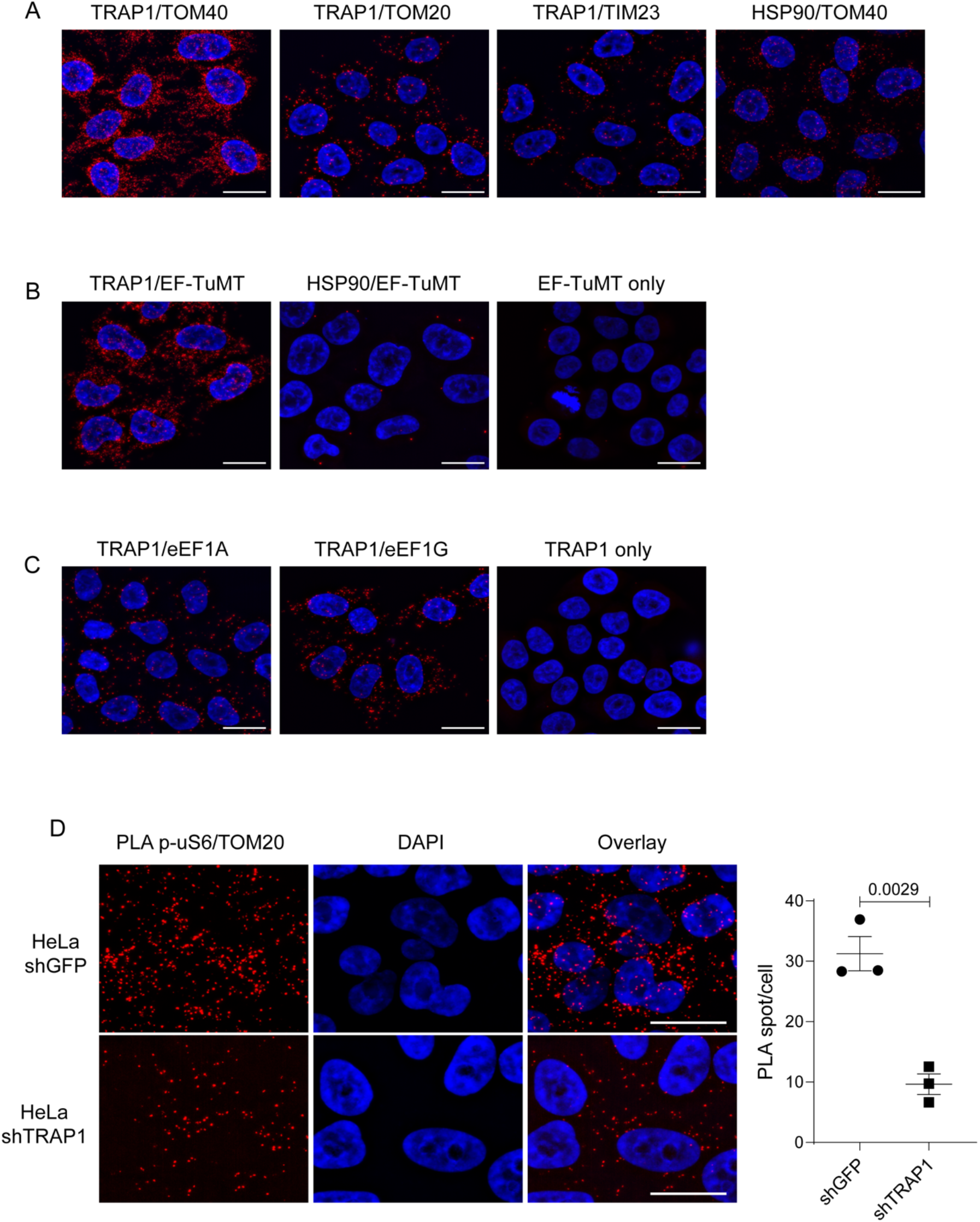
**(A-B-C)** Representative image of PLA showing the interaction, in HeLa cells, between TRAP1 and TOM40, TOM20 and TIM23 (A), between TRAP1 or HSP90 and EF-TuMT (B) and between TRAP1 and EF1A and EF1G. Positive signals of interaction are shown as red dots, nuclei are stained with DAPI (blue). Scale bar = 20 μm. **(D)** Representative image of PLA showing the interaction of TOM20 with phosphorylated (active) ribosomal protein eS6 in HeLa cells following 72hrs induction of TRAP1 and GFP (control)-directed shRNAs. Positive signals of interaction are shown as red dots, nuclei are stained with DAPI (blue). Scale bar = 20 μm. The graph shows the average number of PLA spot/cell, with a p-value representing the statistical significance based on the Student’s t-test (n=3).

